# First near complete haplotype phased genome assembly of River buffalo (*Bubalus bubalis*)

**DOI:** 10.1101/618785

**Authors:** Sudhakar Ananthasayanam, Harish Kothandaraman, Nilesh Nayee, Sujit Saha, Dushyant Singh Baghel, Kishore Gopalakrishnan, Sathish Peddamma, Ram Bahadur Singh, Michael Schatz

## Abstract

This study reports the first haplotype phased reference quality genome assembly of ‘Murrah’ an Indian breed of river buffalo. A mother-father-progeny trio was used for sequencing so that the individual haplotypes could be assembled in the progeny. Parental DNA samples were sequenced on the Illumina platform to generate a total of 274 Gb paired-end data. The progeny DNA sample was sequenced using PacBio long reads and 10x Genomics linked reads at 166x coverage along with 802Gb of optical mapping data. Trio binning based FALCON assembly of each haplotype was scaffolded with 10x Genomics reads and super-scaffolded with BioNano Maps to build reference quality assembly of sire and dam haplotypes of 2.63Gb and 2.64Gb with just 59 and 64 scaffolds and N50 of 81.98Mb and 83.23Mb, respectively. BUSCO single copy core gene set coverage was > 91.25%, and gVolante-CEGMA completeness was >96.14% for both haplotypes. Finally, RaGOO was used to order and build the chromosomal level assembly with 25 scaffolds and N50 of 117.48 Mb (sire haplotype) and 118.51 Mb (dam haplotype). The improved haplotype phased genome assembly of river buffalo may provide valuable resources to discover molecular mechanisms related to milk production and reproduction traits.

## Introduction

Complete genome sequence of a species is essential for studying genetic diversity, inbreeding and breed admixture. Advances in whole genome sequencing (WGS) technologies coupled with genomic data storage and analysing capabilities have helped in utilising the genomic data for genetic improvement, in domestic animals. A fundamental requirement for utilising genomic information is the availability of a reference genome for a species. Complete and accurate reference genome is essential for advanced genomic selection of production traits and gene editing in agriculturally important plant and animal species (Matukumalli et al., 2009). Advances in WGS have enabled sequencing of many species and individuals, but many genome assemblies are left incomplete and fragmented in large part because the underlying sequence reads are too short (Chaisson et al., 2015) and consist of hundreds or even thousands of fragmented contigs and scaffolds. Furthermore, many of these contigs will not be mapped to their chromosomal locations and further enhancing the problem of haplotype variations which are not present in natural populations.

Ideally, an individual’s genome needs to be represented through its different haplotypes, i.e. maternal and paternal haplotypes in diploid organisms. A major goal of a genome project is a complete haplotype-resolved assembly with no gaps (Low et al. 2019). Accurate representation of haplotypes is essential for studies on intraspecific variation, chromosome evolution and allele-specific expression (Koren et al., 2018). Currently available, consensus mosaic genome assemblies do not truly represent the original parental haplotypes. Collapsing haplotypes into a single consensus representation may introduce false variants that are not present in either haplotype, leading to annotation and analysis errors (Korlach et al. 2017). The trio binning, a haplotype phased genome assembly method advanced by Koren et al. (2018) provides an excellent opportunity for increased accuracy of genomic selection in domestic animals. The recent development and application of long-read and linked-read sequencing technologies like PacBio sequel and 10X genomics Chromium have shown considerable promise in improving eukaryotic genome assemblies. Numerous scaffolding technologies have been developed for ordering and orienting assembly contigs, including chromosome interaction mapping (Hi-C) and optical mapping, which provide relatively inexpensive and high-resolution scaffolding data (Bickhart et al. 2017)

Errors in a reference genome assembly can often result in misinterpretations of the underlying sequence of an animal, mainly when the Structural Variants (SVs), such as deletions, insertions, inversions, and more complex rearrangements are the focus for detection. The relatively lower quality of reference genomes produced for livestock species substantially increases the number of false positives generated in the discovery of SVs (Bickhart and Liu, 2014). Even if reads contain a 1% variant-error rate, the combination of eight identical reads that cover the location of the variant will produce a strongly supported variant call with an associated error rate of 10^−16^ (Schatz et al. 2010). Although SV is a form of natural polymorphism, specific or excessive aberrations have been linked to numerous human diseases (Weischenfeldt et al. 2013). Copy number variations (CNV) are a subgroup of structural variations including deletions and duplications. For example, a 660-kb deletion was found to be associated with fertility and milk production in Nordic red cattle (Kadri et al. 2014). Identification of long SVs with haplotype phased genomes will aid in revealing alleles associated with phenotypes, which were hitherto challenging to identify and use them for selection of favoured phenotypes or selection against unfavoured phenotypes (e.g. recessive lethal alleles).

River buffalo (*Bubalus bubalis*) is a very popular domestic animal for milk production in Asia. The estimated population of world buffalo is 224.4 million, of which 219 million (97.58%) are in Asia. India has 113.3 million buffaloes, and they comprise approximately 50.5 per cent of the total world buffalo population (FAOSTAT, 2019). Buffaloes are more resistant to ticks and certain diseases. In comparison to cattle, Buffalo milk contain high fat%. Buffaloes being most widely reared in the developing countries for milk production, developing a reference genome of buffalo would help in the genomic selection of buffaloes for enhancing productivity per animal.

When the present work was initiated, few highly fragmented genome assemblies of Mediterranean, Jaffarabadi and Egyptian buffaloes were available in public domain and river buffalo pseudo sequence aligned to a cattle reference was reported (Tantia et al. 2011) but a good quality genome assembly of *Bubalus bubalis* (river buffalo) was not available and river buffalo was considered as a non-model organism lacking genomic resources such as complete reference genome or annotated gene models (WhitAcre et al. 2017). Currently, a 58% haplotype resolved genome assembly (UOA_WB_1) is publicly available with Genbank assembly accession number (GCA_003121395.1). In the present work, we describe the development of haplotype phased highly contiguous near complete genome assembly of *Bubalus bubalis* by separating haplotypes prior to assembly using a father-mother-offspring trio to accurately and completely reconstruct parental haplotypes.

## Results

Haplotype phased-out genome assembly of the Indian “Murrah” buffalo with an estimated genome size of 2.66Gb required genome sequence data of a trio with a female progeny and her parents. The chosen animal’s genome was sequenced on multiple platforms to capture the complementarity of the sequencing protocols and to span long repetitive regions. We sequenced a total 274Gb on the Illumina platform for the parental samples. On the PacBio sequel, a total of 217.30Gb sequence data was generated of which 186.2Gb was available in reads over 10Kb length with an N50 of 20.46Kb and an average length of 12.85Kb. The 10X Chromium Linked-Read sequencing generated 226.38Gb of the progeny with Long Ranger estimating the molecule length to be 36.91Kb. BioNano Saphyr Chip-based Optical Mapping using the enzyme DLE-1 resulted in 802Gb data with molecules greater than 20Kb. Details on data generated for genome assembly are presented in Table 1. Raw de novo genomic data and Haplotypes have been submitted in NCBI (Bioproject id: PRJNA525182)

**Table 1:** Data Generated to assemble Murrah Genome

| **Datasets generated** | **Total data volume (Gb)** | **Filtered data volume (Gb)** | **QC Filters applied to validate data** | **Data used for the Assembly** |
| --- | --- | --- | --- | --- |
| PacBio Sequel long reads | 217.30 | 186.20 | Reads greater than 10Kb | Reads greater than 500bp |
| 10 X Genomics linked reads | 226.38 | - | None | All |
| Illumina Paired End reads - Parental | 274 | 264 | >Q30 | >Q30 |
| BioNano Optical molecules | 802.72 | 364.94 | >150Kb with 9 labelled sites | >150Kb with 9 labelled sites |

### Trio-Binning based haplotype reconstruction of the Buffalo genome

The short-insert (350bp) Illumina pair-end data of parental samples for the chosen animal were sequenced to generate 274Gb data, to perform a Trio-Binning based genome assembly (Koren et al., 2018) to create a complete diploid reconstruction of the assembly. The PacBio Long-read segregation, for each parental haplotype, was over 99.99%, with less than 140Mb of raw reads being present in the non-classified set, confirming the trio.

The binned PacBio sequel Long-Reads were assembled using both FALCON and Canu, although FALCON produced more contiguous assemblies that we analyzed here. For the Dam haplotype, the assembly length was 2.62Gb in 1837 contigs with N50 of 5.23Mb, and for Sire haplotype the assembly length was 2.62Gb in 1132 contigs with N50 of 9.5Mb. The reconstructed haplotypes were iteratively scaffolded twice using 10X Chromium library generated linked-reads to improve the continuity of the assembly further). The iterative Scaff10x scaffolding improved the assemblies for the Dam haplotype by increasing the assembly length to 2.63Gb in 1130 scaffolds, N50 to 12.44Mb; similarly, the Sire haplotype’s assembly increased to 2.64Gb in 668 scaffolds with an N50 of 40.70Mb. Results are presented in Table 2.

**Table 2:** Consolidated genome assembly metrics of the Murrah Sire and Dam Haplotypes

| Murrah haplotype | Assembly | Software | Assembly level | Number of sequences | Number of gaps | N50 | N90 | Assembly size (Gb) | Max. length(Mb) |
| --- | --- | --- | --- | --- | --- | --- | --- | --- | --- |
| Dam haplotype | PacBio FALCON | FALCON | Contig | 1837 | 0 | 5.23Mb | 1.24Mb | 2.62 | 35.05 |
|  | PacBio Canu | Canu | Contig | 3445 | 0 | 4Mb | 816.17Kb | 2.67 | 21.1 |
|  | PacBio (FALCON)+ 10 X | Scaff10X | Scaffold | 1130 | 70.7Kb | 12.41Mb | 3.18Mb | 2.63 | 40.11 |
|  | PacBio (FALCON)+ 10 X + Optical mapping | BioNano Access | Scaffold | 64 | 33.74Mb | 83.23Mb | 42.16Mb | 2.64 | 156.98 |
| Sire haplotype | PacBio FALCON | FALCON | Contig | 1132 | 0 | 9.5Mb | 2.43Mb | 2.62 | 38.34 |
|  | PacBio Canu | Canu | Contig | 2883 | 0 | 6.21Mb | 1.32Mb | 2.69 | 35.05 |
|  | PacBio (FALCON) + 10 X | Scaff10X | Scaffold | 668 | 48.9Kb | 42.70Mb | 9.1Mb | 2.64 | 104.72 |
|  | PacBio (FALCON) + 10 X + Optical mapping | BioNano Access | Scaffold | 59 | 17.44Mb | 81.98Mb | 42.24Mb | 2.63 | 156.63 |

### Optical Map of the Buffalo genome

The BioNano Saphyr molecules were filtered to retain molecules with a length greater than 150Kbp and nine nicking sites, leading to a final coverage of 137.20x hosting 364.94 Gb of the original dataset. The Bionano optical molecules assemblage produced 340 genome maps with an N50 of 75.85Mb. Filtered molecules were mapped back to the genome map showing the confidence of 29.2 with 80% alignments (Table 3).

**Table 3:**
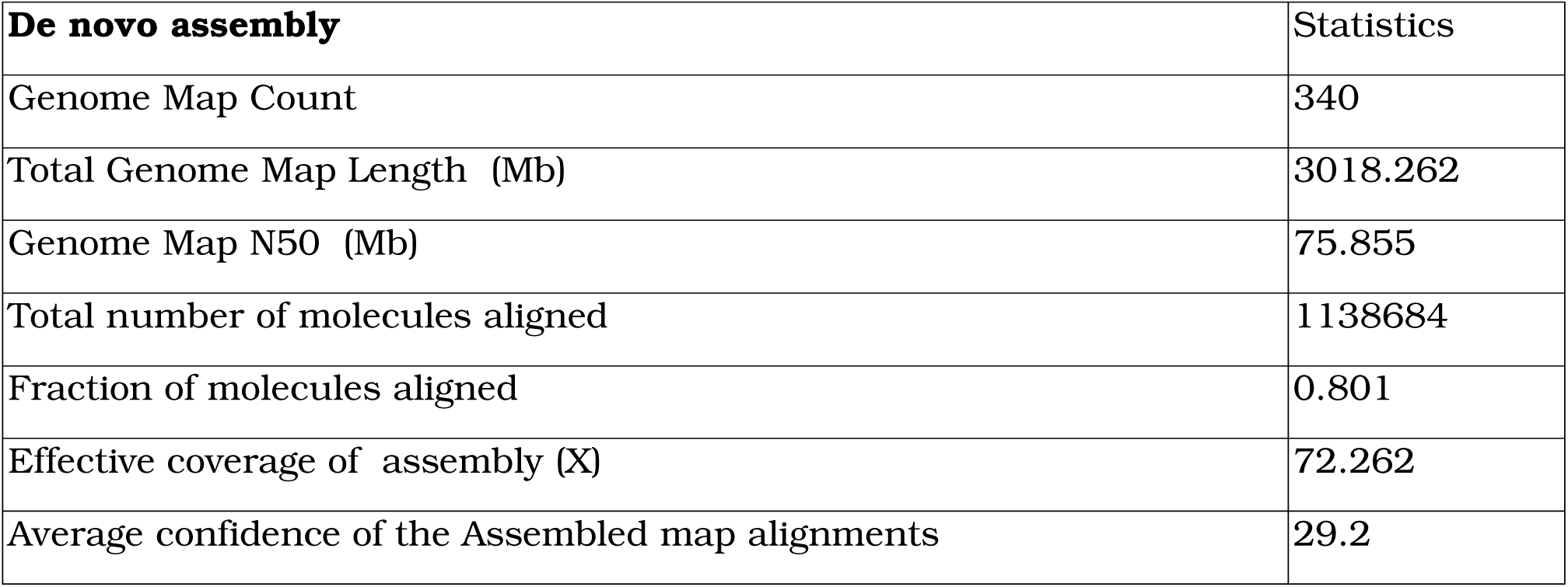
Optical map assembly metrics of the Buffalo genome

The assembled 340 genome maps were aligned against the recently published Mediterranean buffalo, highlighting an alignment of 2317.902Mb, with an alignment of 87.3% over the whole genome. Notably, the optical map assembly was able to capture the entire chromosome arm for chromosome 16 (**Figure 1**).

**Figure 1:**
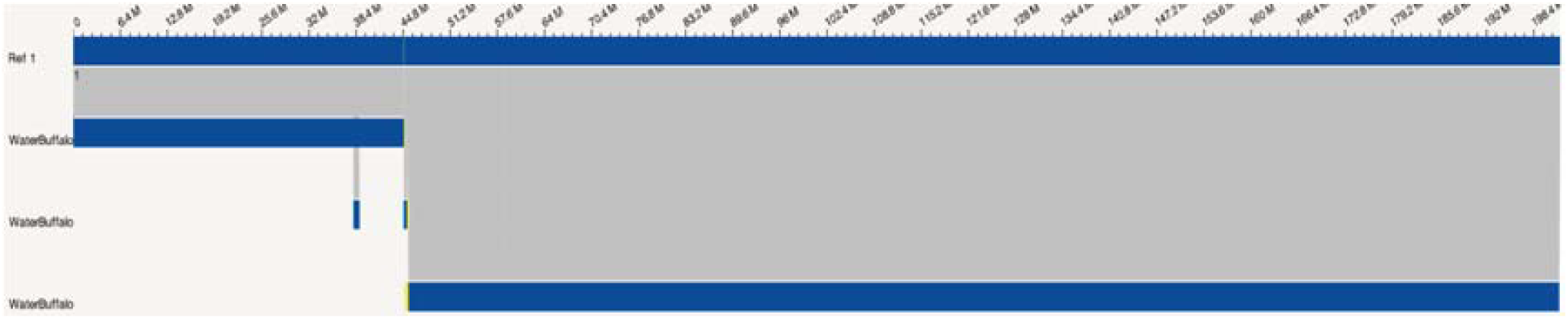
BioNano Access snapshot showing the complete assembly of the chromosomal arm for the chromosome 16 against reference Mediterranean River buffalo

The BioNano molecule assembly was used for hybrid scaffolding of the two selected haplotype assemblies. Scaffolded sequences reported by BioNano Access contained greater than 2.64 Gb (Supplementary Table 1) of the estimated 2.66Gb genome, leading to a genome completion of 99.20%.

The hybrid scaffolding of the FALCON+10X scaffolds with BioNano genome maps produced 64 and 59 scaffolded sequences representing over 99% of the Dam and Sire haplotypes, with only 27.1Mb and 19.98 Mb remaining unscaffolded represented in over 640 and 480 sequences. The Dam and Sire Haplotype scaffolded assemblies contained 33.74 Mb and 17.44Mb “N”s respectively. Haplotype coverage against latest available reference genomes is reported in Table 4.

**Table 4:**
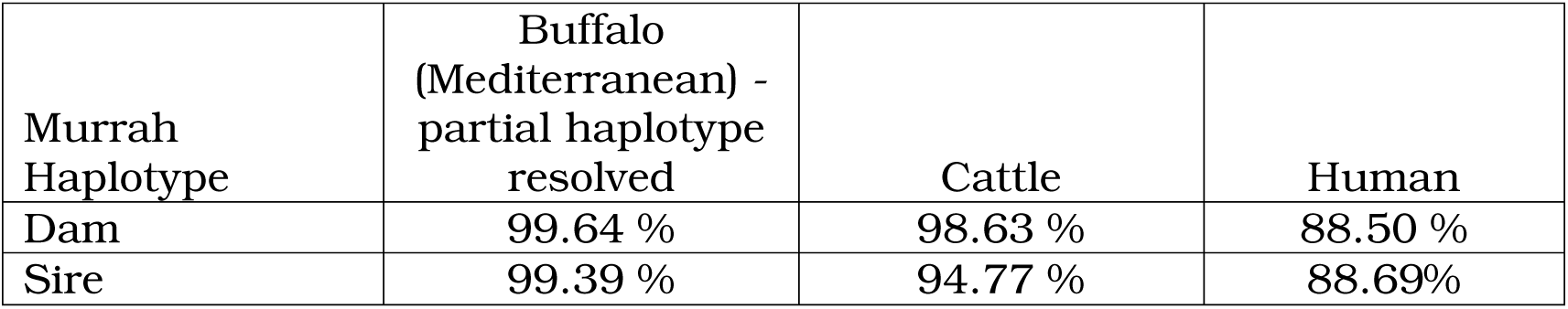
Haplotype coverage of Murrah buffalo against latest available haplotype phased genomes

| Murrah Haplotype | Buffalo (Mediterranean) - partial haplotype resolved | Cattle | Human |
| --- | --- | --- | --- |
| Dam | 99.64 % | 98.63 % | 88.50 % |
| Sire | 99.39 % | 94.77 % | 88.69% |

Benchmarking universal single copy orthologs (BUSCO) (Simao et al. 2015) analyses revealed 91.25% and 92.06% complete universal single copy orthologs on Hybrid-scaffolded assemblies, respectively of maternal and paternal haplotypes on the mammalian lineage consistent with Mediterranean buffalo (Low et al. 2019). Similarly, BUSCO analyses on the Vertebrata lineage revealed 93.8% and 94.4% for the maternal and paternal haplotypes respectively. BUSCO analyses on the chosen polished optical map based hybrid scaffolding (Supplementary Figure 1).

### Structural variant discovery

#### Assemblytics based structural variation identification

We identified structural variants in the assembled haplotypes using Assemblytics (Nattestad et al. 2016) relative to several related species. Briefly, Assemblytics identifies structural variations as breaks within or between alignments computed by the whole genome alignment algorithm MUMmer. To broadly catalog structural variations, we considered several related species including: the *Bos frontalis* (available from GigaScience, BioProject: PRJNA387130, *Capra hircus* (GCA_001704415.1), *Bos indicus* X *Bos taurus* Crossbred (GCA_002263795.2), *Homo sapiens* (GCA_000001405.27, GRCh38, primary assembly) and the recently published Mediterranean buffalo (*Bubalus bubalis*, GCA_003121395.1). This analysis revealed a plethora of deletion, repeat contraction and expansion events.

Comparison with the other parental haplotype, i.e., the dam haplotype showed that approximately 19.34Mb represented the genomic variations.

The Murrah Sire Haplotype (SH) Assemblytics comparison with chosen related species are summarised below in **Table 4a,b**. Log scaled distribution of structural variants is visualised in **Figure 3**.

**Figure 3:**
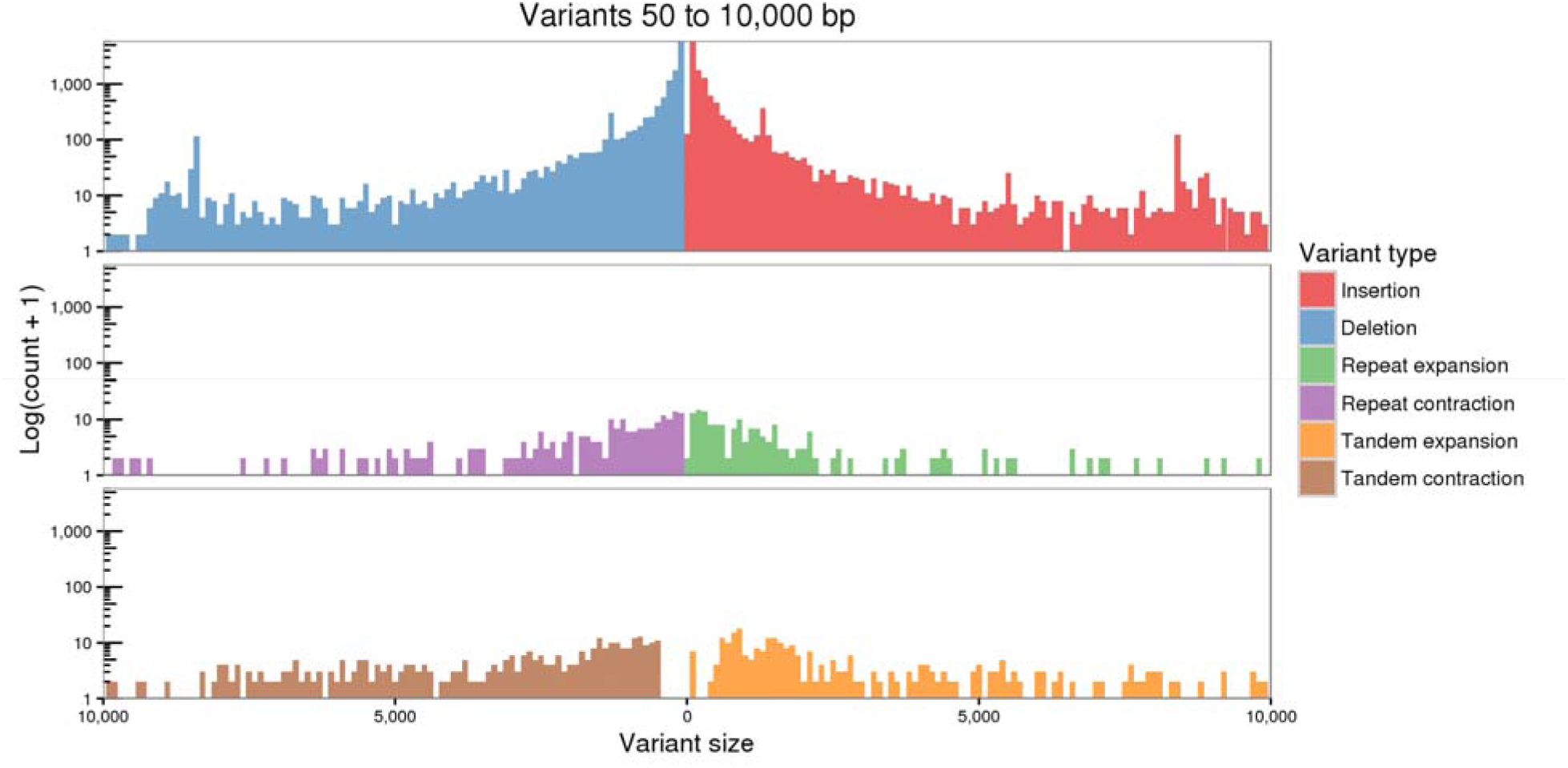
log-scaled distribution of variants. Major structural variant event classes were insertions and deletions, especially in the 500-10000bp range. A total of 835 Contraction and Expansion events contributed to nearly 2Mb of the structural variant length.

**Table 4a:**
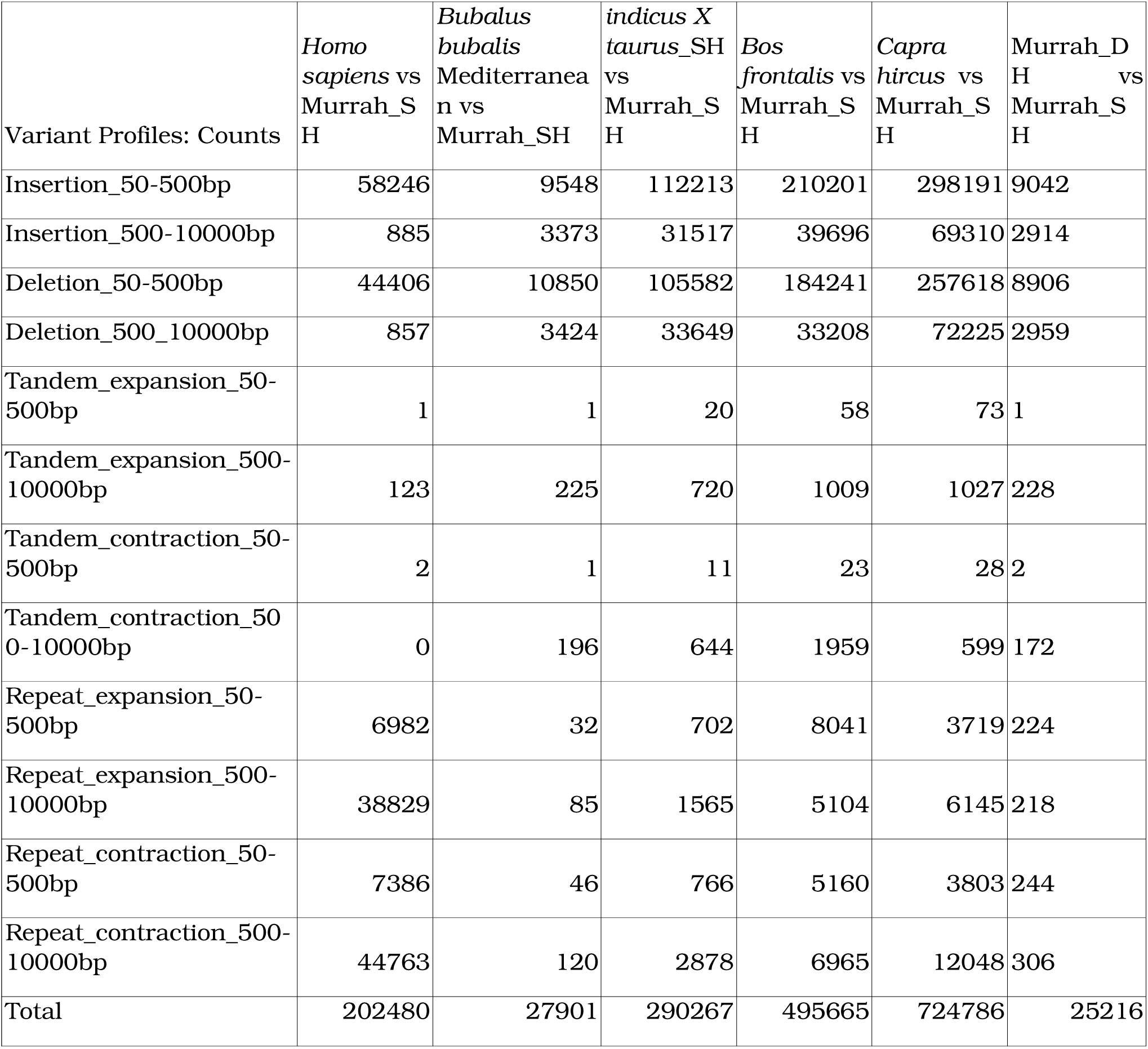
Variant counts of 50-500bp and 500-10000bp as inferred by Assemblytics:

**Table 4b:** Cumulative variant lengths of 50-500bp and 500-10000bp as inferred by Assemblytics:

| Variant Profiles: Length | *Homo sapiens* vs Murrah_S H | *Bubalus bubalis* Mosaic vs Murrah_S H | *Bos indicus* X *Bos taurus*_SH vs Murrah_S H | *Bos frontalis* vs Murrah_S H | *Capra hircus* vs Murrah_S H | Murrah_D H vs Murrah_SH |
| --- | --- | --- | --- | --- | --- | --- |
| Insertion_50-500bp | 9586539 | 1605227 | 21533259 | 32668870 | 62811577 | 1441292 |
| Insertion_500-10000bp | 1061391 | 7487722 | 54268134 | 68306084 | 98561308 | 6674070 |
| Deletion_50-500bp | 7741268 | 1725212 | 20596636 | 31563303 | 52865775 | 1434394 |
| Deletion_500_10000bp | 956777 | 7761808 | 56591287 | 46473160 | 107104864 | 7119507 |
| Tandem_expansion_50-500bp | 103 | 437 | 4778 | 14015 | 20300 | 147 |
| Tandem_expansion_500-10000bp | 744492 | 672085 | 1729372 | 2284749 | 2322628 | 826763 |
| Tandem_contraction_50-500bp | 351 | 492 | 3095 | 7350 | 5936 | 321 |
| Tandem_contraction_500-10000bp | 0 | 631634 | 1630833 | 2919783 | 1376429 | 491241 |
| Repeat_expansion_50-500bp | 1867468 | 7926 | 173631 | 1473246 | 932438 | 48505 |
| Repeat_expansion_500-10000bp | 107679795 | 167762 | 2982070 | 7719070 | 10047730 | 520148 |
| Repeat_contraction_50-500bp | 1983849 | 11094 | 192563 | 1188840 | 956065 | 54375 |
| Repeat_contraction_500-10000bp | 136724088 | 314172 | 6611224 | 14360739 | 27975575 | 739169 |
| Total | 268346121 | 20385571 | 166316882 | 208979209 | 364980625 | 19349932 |

#### Pseudo-Molecule Construction

The sire and dam haplotypes were aligned to the Mediterranean buffalo genome assembly’s chromosomal sequences using MiniMap2 (Li, 2018). RaGOO assigns unmapped sequences, i.e. sequences that had no alignments with the reference as “Chr0_contig” padded with “N”s. We did not observe any such sequences in case of the sire haplotype, highlighting the quality of the haplotype resolved assembly, as well as no ambiguities when considered with the reference. The dam haplotype had approximately 5.24Mb unlocalized sequences, concatenated and placed in “Chr0_RaGOO”.

Finally, RaGOO was used to order and build the chromosomal level assembly with 25 scaffolds and N50 of 117.48 Mb (sire haplotype) and 118.51 Mb (dam haplotype).

Comparison of the RaGOO ordered chromosomal sequences by treating the Sire Haplotype as the reference and dam haplotype as a query (including the unlocalized sequences), led to the identification of 26,489 variants with a cumulative length of 23.06Mbp. Major classes of these variants were insertion and deletions events between 500-10000bp with almost 6.7Mb in haplotypes.

The alignments were further sub classified, observing 9241 unique alignments, and 2214 repetitive alignments between the Dam and the Sire haplotype assemblies. Finally, alignments over 50kb in length were then visualised using the R package Circlize to plot chord diagrams (Gu,2014).

**Figure 4:**
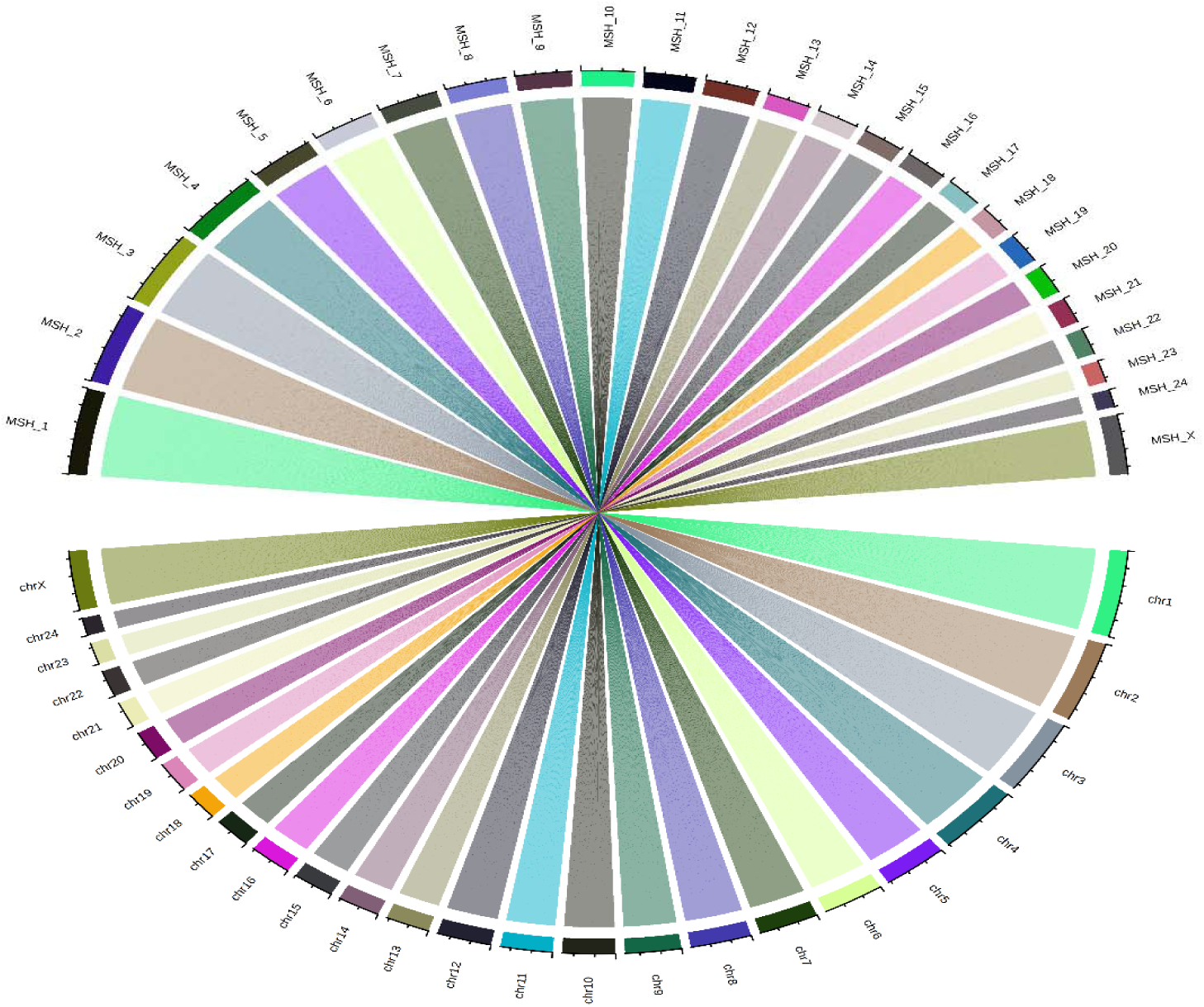
Alignments over 50Kb in length between the Murrah and Mediterranean buffalo assemblies plotted as a chord diagram. Murrah Sire haplotypes chromosomes referred as MSH_”x” and Mediterranean buffalo chromosomes as “chr n”, where x was the chromosomal assignment based on RaGOO.

**Figure 5:**
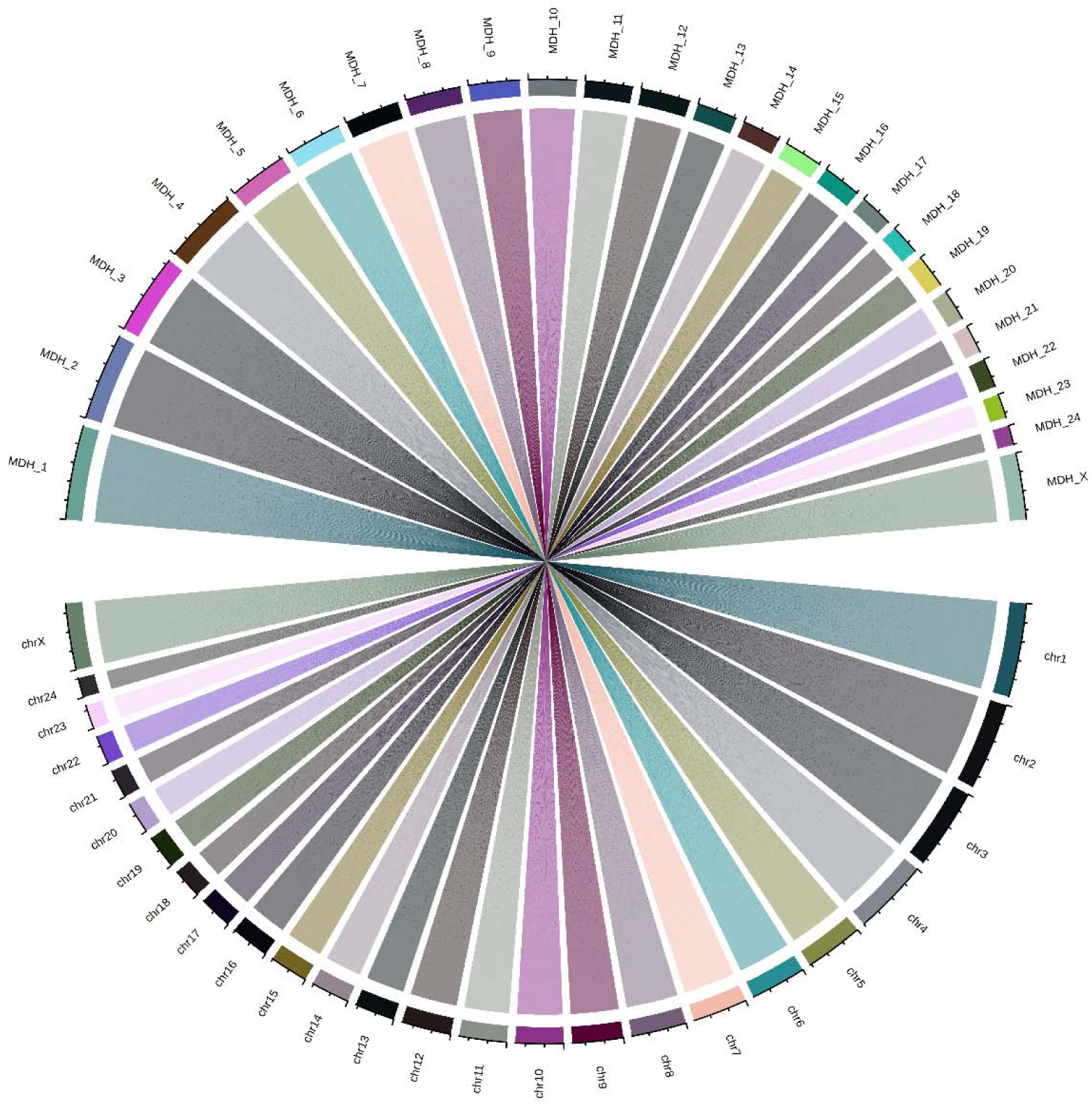
Alignments over 50Kb in length were plotted in the chord diagram. Murrah Dam haplotypes chromosomes referred as MDH_”x” and Mediterranean buffalo chromosomes as “chr n”, where x was the chromosomal assignment based on RaGOO

## Methods

### Reference Animal selection and sample collection

A true to type Murrah female calf maintained in Government livestock farm, Hissar, Haryana was used as a candidate for de novo genome assembly. The dam was available at same farm, while the sire of the progeny was reared at The Sperm station, Hissar. Pedigree of the de novo candidate was confirmed with 13 microsatellite markers, and Karyotype was identified as normal. Blood samples of all the three animals were collected in EDTA vacutainers as per Institutional animal ethics committee guidelines.

### Genomic DNA isolation, Sequencing and Optical mapping

The high molecular weight (HMW) DNA was isolated from blood samples using QIAGEN MagAttract HMW DNA kit (Cat No: 67563) for PacBio Sequel, Chromium 10X libraries and Paired-End libraries. BioNano kit – Blood and Cell Culture DNA isolation kit and modified protocol developed by Nucleome Informatics Pvt. Ltd. (Cat No. 80004) were used for extracting the DNA required for the Saphyr based Optical molecule generation. DNA isolation and quality checks were performed as per manufacturer’s recommendations.

PacBio libraries were prepared with SMRTbell Express Template Preparation Kit with 5μg gDNA of 30 – 40 kb size. Bluepippin size selection was performed at 15-50 kb. 10X Genomics Chromium libraries were prepared using Chromium™ Genome HT Library Kit & Gel Bead Kit v2, 96 reactions (Cat No. PN-120261). Latest Illumina TruSeq Strand-specific PCR free Library kit (Cat. No. 20015963) was used for 350 bp insert size library preparation. All libraries were prepared as per the manufacturer’s protocol.

Sequencing: The DNA libraries were sequenced on a PacBio sequel system v2 with Sequel v4 Kit. 10X Genomics Chromium libraries and Illumina Truseq libraries were sequenced in Illumina NovaSeq 6000 platform.

### Optical Mapping

HMW DNA was extracted with Bionano kit – Blood and Cell Culture DNA isolation kit (Cat No. 80004) for the Saphyr based Optical Mapping. For Bionano sample preparation, Quick-thawed frozen sample aliquot at 37°C were used and WBCs were embedded in agarose plugs; plugs were treated with Proteinase K to extract DNA from plugs. Agarose treatment was performed for DNA recovery from Agarose plugs and drop dialysis was performed to extract clean DNA. The DNA concentration was measured using the PicoGreen assay. DLE-1 labelling of DNA was done at specific sequence motifs with Direct Label and Stain (DLS) chemistry, a direct labelling enzyme attaching a fluorophore (green) directly to the DNA at a 6-mer recognition site. Optical molecules were generated with BioNano Saphyr Chip as per manufacturer’s protocols.

### De Novo Genome Assembly

#### Trio-Binning and Long Read Assembly

Initially, the parental Illumina data was screened for adapters, low quality and ambiguous bases using Fastp, and data above Q30 were retained for subsequent Trio-Kmer generation.

The genome assembly was approached using the PacBio reads at a primary assembly level using a trio-based binning method using the parental haplotypes as per Koren et al. 2018. Initially the data was partitioned at a kmer of 21 and then subsequently at 18 considering a minimum kmer coverage of 10 using Meryl.

The reads belonging to each of the haplotype-specific bins were assembled separately to generate a diploid assembled genome, using FALCON and Canu assemblers (Chin et al. 2016 and Koren et al. 2017). We polished the assemblies by Arrow from SMRTLink 6.0 using Haplotype specific binned reads. We chose the FALCON assemblies for further scaffolding based on the superior contiguity metrics.

#### 10X Chromium Scaffolding

10X Genomics linked reads were initially debarcoded as per the recommendations of Scaff10Xv3, and BWA-MEM was used to map the renamed 10X chromium reads to derive a SAM file. Scaff10Xv3 then uses a relational matrix to identify the links between the contigs based on the 10X barcodes, which is then used to join the scaffolds. The assemblies were iteratively scaffolded twice using Scaff10X (Table 2).

#### BioNano map generation and hybrid scaffolding

The BioNano^®^ molecules were filtered to retain molecules over 150Kb with a minimum of 9 DLE-1 nicking endonuclease-specific recognition sites. This data was assembled using BioNano Access software, using the De Novo Assembly suite to perform the assembly (Table 3). The assembly was retrieved in a CMAP format to subsequent using the maps for a scaffolding step.

Further scaffolding of the genome was performed using the assembled BioNano genome Maps using a single enzyme workflow process (DLE-1) having a recognition site CTTAAG. The scaffolded sequences were finally polished using the debarcoded 10X reads using Pilon version 1.22.

The resulting assemblies were labelled as Murrah_DH (MDH) and Murrah_SH (MDH) to signify the assemblies were arising from the haplotypes and had been phased.

#### Genome completeness study

All the subsequent analyses were performed on the polished genome assemblies. BUSCO analysis was performed on the chosen FALCON assemblies and optical map based hybrid scaffolded assemblies at the vertebrata and mammalia_odb9 lineage-specific profile at default parameters to assess the quality of the assemblies. gVolante (Nishimura, 2017) Core Eukaryotic Genes Mapping Approach (CEGMA) (Yuichiro et al. 2015) was performed to assess the conservancy of Core Vertebrate Genes.

#### Chromosomal ordering of scaffolds

The BioNano scaffolded and polished sequences were ordered into chromosomes using RaGOO by breaking chimeric scaffolds (Alonge et al. 2019) using only the chromosomal sequences from the Mediterranean river buffalo genome (Low et al. 2019).

### Genome-wide structural variations discovery

The final scaffolded assembly from the Sire haplotype was chosen for comparison with the recently published Mediterranean water buffalo genome (Low et al. 2019), *Bos frontalis* (Gayal) (Wang, et al. 2017), *Capra hircus* ARS1 (goat), *Homo sapiens*. In each case, we retained the “chromosomal” sequences, except in the case of the *Bos frontalis* genome which was a scaffold level assembly in over 460000 scaffolds.

We treated the Sire haplotype, containing 59 scaffolds as the query and the above genomes as a reference including the dam haplotype assembly, which was then aligned using Minimap2 (Li et al. 2018) at default parameters. The output SAM file was then parsed using “sam2delta.py” script from RaGOO, to retrieve the alignments in a NUCmer compatible delta format. The delta file was submitted to Assemblytics (Nattestad et al. 2016) to identify structural variants from 50bp-10Kbp.

## Discussion

Genomic selection in dairy cattle has led to rapid genetic gains per generation. Long-range linkage disequilibrium (LD) may be extensive in the reference population animals, causing large chromosomal segments or haplotypes to be common. Consequently, there will be many combinations of SNPs that explain the effect of the haplotype as well as the causal mutations (Meuwissen et al. 2016). High quality haplotype resolved genomes will aid in direct identification of causal complex structural variants.

During the earlier years of the NGS revolution classically Genomes were chiefly assembled using short read approaches. However, short read based genome survey analyses can play a significant role in deciding the strategy for genome assemblies and subsequent data analyses (Reddy et al. 2018.). The major fallacies of using short read approaches include being unable to span highly repetitive regions longer than the read length, haplotype collapsing or heterozygosity resolution. The long read sequencing aims to resolve the above issues, by being able to span the repeats by leveraging read length running in kilobases but with significantly more errors, using a mosaic approach wherein the parental haplotypes are collapsed while assembling the genome resulting in calling of false-positive or erroneous alleles, poor annotations and downstream analyses.

Recent methodologies incorporate algorithms that can assemble, classify contigs and alternative contigs. However, these genomes might still incorporate phase switching errors depending on the resolution of the assembly graph (Koren et al. 2018). To resolve this, in case of the Indian Murrah buffalo Trio-Binning approach has been used which uses the k-mers from the parental Illumina PE datasets to partition and bin the long-read data of the progeny into haplotype-specific bins, later assembled using FALCON and Canu. The Canu assembly was more fragmented; hence FALCON assembly was taken up for further scaffolding.

Use of 10X Chromium datasets brought a twofold reduction in the number of contigs by initially consuming the datasets to scaffold the FALCON assemblies with a minimal intrusion of ambiguous bases, but improve the contiguity. Iterative scaffolding reduces the number of scaffolds by strengthening the links associated with the 10X dataset and reordering the contigs incorporated as needed.

BioNano Saphyr based Optical Mapping has been demonstrated as a powerful tool for correcting, orienting and reordering the NGS based contigs. Preparation of the Ultra High Molecular Weight gDNA is critical in achieving a high-quality, comprehensive genome map. Highly stringent filters to retain molecules beyond 150Kb length, nine labelling sites and a good signal/noise ratio are essential to building highly contiguous DeNovo genome maps. In this study, Super-scaffolding the 10X assisted FALCON scaffolds of the haplotypes using the BioNano DeNovo Genome maps gave a significant reduction (> 11X) in the number of scaffolds in each of the haplotypes. Haplotype coverage of domesticated animals against latest available reference genomes and review of published research papers indicate that Murrah haplotypes have best haplotype resolution among domesticated animals.

Chromosomal length assemblies can be achieved by using Chromatin conformation capture technologies like Hi-C to build upon the pre-existing assembly or scaffolds. Alternately, RaGOO can be used to order the contigs/scaffolds into chromosomes in case a reference genome is already available to be used as a genetic map, which can be an economical method to build a chromosomal scale assembly and validation of the scaffolds in assembly. Hi-C can be used as an independent evaluation to validate the scaffold links generated by RaGOO. However, Hi-C scaffolds can also have inversion errors in case of highly fragmented assemblies, which can be overcome if the assembly graph is known.

RaGOO based chromosomal assembly of the Indian Murrah haplotypes led to the identification of the structural variants using Assemblytics and genomic comparison. Currently, the Hi-C data from the same Murrah female animal is being generated to scaffold the finalised BioNano map aided genome scaffolds to improve genome assembly quality and compare the ordering by RaGOO. Complete transcriptome information of Bubalus bubalis would provide a valuable resource to infer exon-intron boundaries and study functional genomic regions by annotating the genes.

With present experience in triobinning, we recommend the use of Optical mapping for super scaffolding on long PacBio reads and 10X chromium synthetic linked reads for developing a highly contiguous genome assembly.

The newly developed high-quality haplotype phased buffalo genome assembly shall provide genomic resources for identification of trait specific structural variants, development of SNP array chip for genotyping and augmenting genomic selection procedures to improve milk production and reproduction traits of buffaloes in developing countries.

## Supplementary data

All supplementary data are provided below.

## URLs

http://www.fao.org/faostat/en/#data/QL downloaded on 16 March 2019. Scaff10x: https://github.com/wtsi-hpag/Scaff10X

## ACKNOWLEDGEMENTS

The authors humbly thank The International Development Association (IDA) of World Bank and Shri Dilip Rath, Mission Director, National Dairy plan (NDP) for funding and necessary support. We also thank Dr C G Joshi, Director, Gujarat Biotechnology Research Centre, Gandhinagar for technical help, Dr Om Girdhar Singh (HLDB), Dr Satpal and staff of Government livestock farm, Hissar and The Sperm station, Hissar for providing samples as required.

We also thank NDDB management for funding and supporting liberally for this project.

We would like to acknowledge Dr. Sara Goodwin from Cold Spring Harbor Laboratory and Chai Fungtammasan from DNANexus for their technical guidance during sequencing and analyses.

## Disclaimer

Mention of trade names or commercial products or institutions in this article is solely for providing accurate information and does not imply any recommendation by National Dairy Development Board, India or its subsidiaries.

---

**Supplementary Table 1:** Hybrid Scaffolding using BioNano Optical Maps.

|  | Scaffolded Sequences |  |
| --- | --- | --- |
| Optical Map Assisted Scaffolding | Murrah_Dam | Murrah_Sire |
| NG50 length | 76022339 | 73558747 |
| NG50 length (Mb) | 76.02Mb | 73.55Mb |
| L90 size | 29 | 29 |
| Total number of bases | 2646160752 | 2639523255 |
| Total number of bases(Gb) | 2.65Gb | 2.64Gb |
| N50 length | 83232536 | 81982981 |
| N50 length (Mb) | 83.23Mb | 81.98Mb |
| Number of scaffolds | 64 | 59 |
| Maximum length | 156982567 | 156633660 |
| Maximum length (Mb) | 156.98Mb | 156.63Mb |
| N90 length | 42161065 | 42242199 |
| N90 length (Mb) | 41.16Mb | 42.24Mb |
| Gaps in Assembly | 33709104 | 17432725 |
| Gaps in Assembly (Mb) | 33.70Mb | 17.43Mb |
|  | Unscaffolded Sequences |  |
| Total number of bases | 27129257 | 19983335 |
| Total number of bases(Mb) | 27.12Mb | 19.98Mb |
| N50 length | 50733 | 46338 |
| N50 length (Kb) | 50.73Kb | 46.33Kb |
| Number of scaffolds | 641 | 482 |
| Maximum length | 776203 | 781035 |
| Maximum length (Mb) | 7.76Mb | 7.81Mb |
| Gaps in Assembly | 1300 | 1000 |
| Gaps in Assembly (Kb) | 1.3Kb | 1Kb |

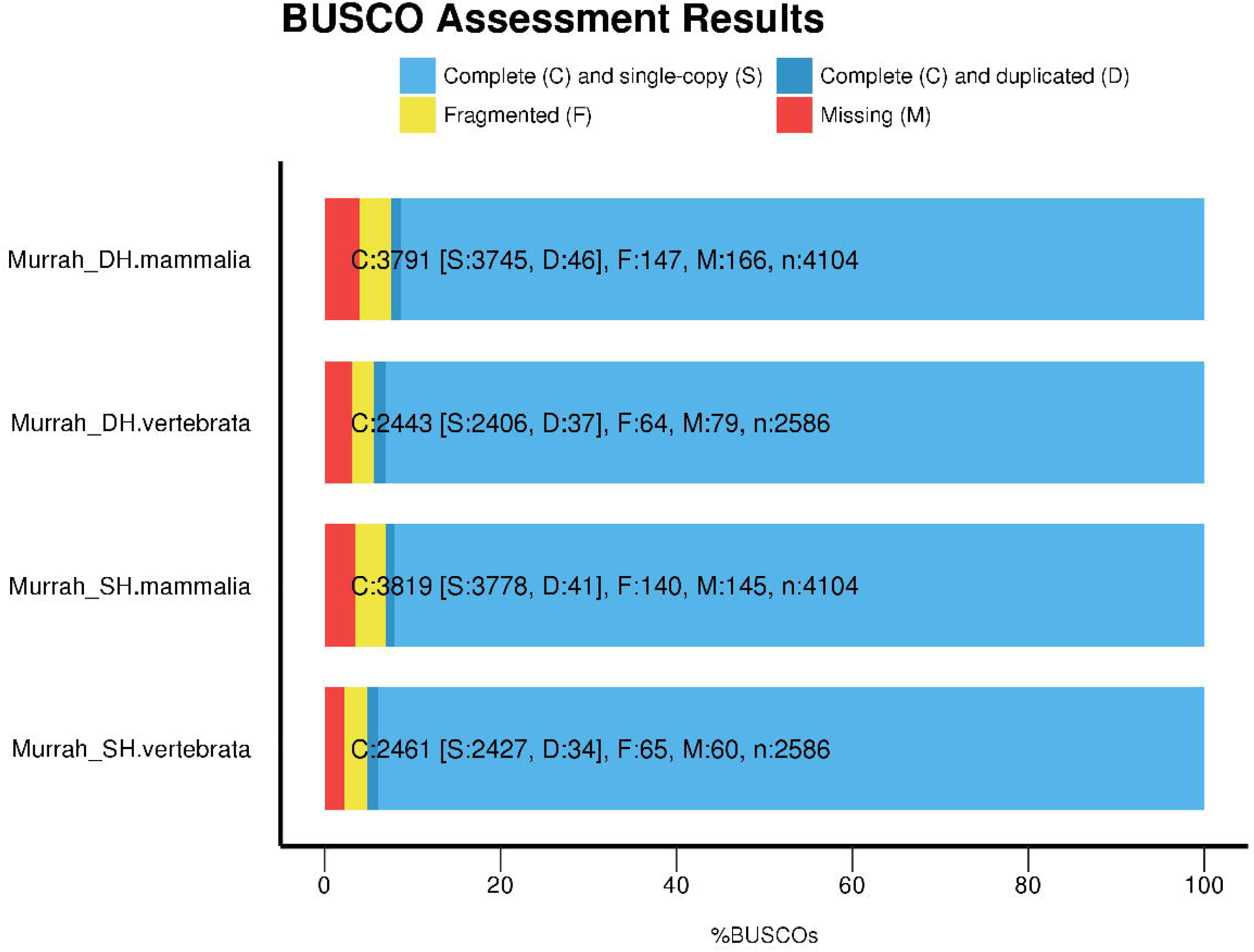

